# PRMT activity promotes global 3’ UTR shortening in proliferating cells

**DOI:** 10.1101/2025.03.06.641848

**Authors:** Llywelyn Griffith, Charlotte Capitanchik, Shaun Moore, Anca Farcas, Mags Gwynne, Martina Pedna, D. Marc Jones, Anob M. Chakrabarti, Dimitris Lagos, Jelena Urosevic, James T Lynch, Jernej Ule

**Affiliations:** UK Dementia Research Institute at King’s College London, London, UK; The Francis Crick Institute, London, UK; Research and Early Development, Oncology R&D, AstraZeneca, Cambridge, UK; Hull York Medical School, University of York, UK; UCL Respiratory, Division of Medicine, University College London, London, UK

## Abstract

Protein methyltransferase (PRMT)-catalysed arginine methylation is a widespread post-translational modification that regulates numerous RNA-binding proteins and frequently becomes dysregulated in cancer. While PRMT inhibitors have shown promise as an anti-cancer strategy, greater understanding of the downstream pathways linking arginine methylation to tumour-promoting phenotypes is needed to improve patient stratification and develop more effective therapeutic approaches. Here, we reveal arginine methylation as a critical regulator of alternative polyadenylation (APA) patterns that are fundamental to tumour progression. 3′ RNA-sequencing assays uncover a rapid and global shift toward longer 3′ UTR isoforms upon dual (symmetric and asymmetric) methylation (DMAi), impacting a broad range of cellular proliferation and signalling genes. Arginine methylation is required for sustaining proximal poly(A) site usage under high proliferative demand, as DMAi treatment blocks use of such sites in activated T cells, various cancer cell lines and patient-derived lung organoids. DMAi also counteracts the 3′ UTR shortening caused by reduced CFIM25 expression, which normally promotes oncogenic isoforms. DMAi treatment affects APA in many of the same mRNAs as impaired cleavage and polyadenylation activity, and these mRNAs contain characteristic signatures such as high GC-content and long 3’ UTRs. This systematic impact of PRMT activity on APA regulation broadens the potential utility of PRMT inhibitors as therapeutic agents for both cancer and immune-related diseases.

## INTRODUCTION

Protein methyltransferases (PRMTs) catalyse the addition of methyl groups onto arginine residues within proteins, utilizing S-adenosyl methionine (SAM) as a donor molecule. Arginine methylation alters the hydrophobicity of arginine side chains, impacting protein-protein and protein-RNA interactions, and thereby affecting protein function and localization^1^. Type 1 PRMTs (PRMT1-4, PRMT6 and PRMT8) and Type II PRMTs (PRMT5 and PRMT9) first monomethylate arginines and then catalyse either asymmetric dimethylation (ADMA) or symmetric dimethylation (SDMA), respectively^2^. PRMT1 and PRMT5 are generally the most abundant of these enzymes, and thus primarily responsible for ADMA and SDMA. They target a broad range of substrates including histones, RNA-binding proteins (RBPs) and DNA-damage repair factors, and therefore arginine methylation can directly impact a variety of cellular processes such as gene expression, intracellular signalling and cell cycle progression^1^. However, while the impact of SDMA on alternative splicing has been well explored^3,4^, other RNA-processing mechanisms regulated by PRMT activity across cell types remain poorly understood.

Arginine methylation represents a particularly attractive therapeutic target given the observation that in 10-15% cancers, the gene encoding MTAP - a key component of the methionine salvage pathway - is co-deleted alongside the tumour suppressor gene CDKN2A^5,6^. This deletion leads to an accumulation of methylthioadenosine (MTA), a natural inhibitor of PRMT5, rendering these tumours especially vulnerable to further PRMT disruption and thus expanding the potential therapeutic window for PRMT inhibitors. Moreover, arginine methylation levels are up to 3-fold higher in immortalised and cancer cells compared with primary cells, and are highest in cells isolated from human metastases^7^, indicating that they could promote the cancer-related regulatory programmes. Indeed, increased Type I and Type II PRMT activity has been implicated in numerous cancers, leading to the development and clinical evaluation of several inhibitors targeting both these enzymes^8–16^. Type I and Type II inhibitors exhibit synergistic effects when combined^11^, underscoring the cellular crosstalk between both modes of dimethylation. However, first-generation PRMT5 inhibitors demonstrated limited clinical activity, due to dose-limiting haematological toxicities^17,18^. Greater understanding of the essential roles of PRMTs in normal cells in relation to roles that are particularly important for tumour cells could support further development of targeted therapies and improve patient stratification.

Proper assembly of the spliceosome is dependent on PRMT5-catalysed arginine methylation of its core Sm components^19^, while ADMA and SDMA also regulate the activity of several auxiliary splicing factors such as SRSF1^20^ and HNRNPA1^21^. Indeed, multiple groups have demonstrated disrupted splicing of key cell cycle and DNA-damage repair genes upon PRMT inhibition^3,11,22–24^. A growing number of mass spectrometry studies^3,11,21,25,26^ have also identified RBPs functioning in alternative polyadenylation (APA) and translation as PRMT1 and PRMT5 substrates. Despite the critical importance of APA in cellular proliferation and tumour progression^27^, its modulation by arginine methylation has not yet been thoroughly investigated.

Here, we find that arginine methylation is required for recognition of proximal poly(A) sites that are associated with highly proliferating cells, such as activated T cells or cancer cells. Inhibition of ADMA and SDMA triggers global 3’ UTR lengthening across a broad range of cancer models and in T cells, affecting key genes in a manner that helps to explain the inhibitory effects on cell proliferation.

## RESULTS

### Inhibition of arginine methylation triggers a global switch to distal poly(A) sites

Numerous core and auxiliary cleavage and polyadenylation (CPA) factors, including WDR33, PCF11, CSTF2, and CPSF6, have been identified as possessing either SDMA or ADMA modifications in recent mass spectrometry analyses^3,11,21,25,26^. These proteins form part of a broader CPA machinery that assembles on pre-mRNA at poly(A) sites. Many genes have multiple poly(A) sites, allowing the generation of several transcript isoforms with variable 3’ UTRs through the process of alternative polyadenylation (APA). Notably, T cell activation was found to induce major changes in APA, which lead to global shifts toward proximal poly(A) sites that produce isoforms with shorter 3’ UTRs^28^. Similarly, numerous studies have identified global 3’ UTR shortening as a recurrent feature across many cancer types^28,29^.

Given the arginine methylation of key CPA-related RBPs, we were motivated to assess its impact on APA. We performed 3’ RNA-seq in LU-99 cells - an MTAP-null NSCLC cell line - treated for 96 hours with LLY-283 to inhibit SDMA, GSK-715 to inhibit ADMA, or with both compounds for double inhibition of SDMA and ADMA (Figure S1A). We included double inhibition to prevent cellular compensation mechanisms, given the observation that ADMA inhibition causes many substrates to be scavenged by SDMA methylation^30^ (Figure S1A). To analyse APA, we firstly derived a poly(A) site atlas across all samples as described in Hallegger et al. 2021^31^, followed by DRIMSeq^32^ of differential poly(A) site usage upon methylation inhibition. We categorised genes containing significant APA events according to whether the affected poly(A) sites were located within the same terminal exon (TUTR) or alternate terminal exons (ALE) (Figure S1B). Moreover, genes were termed as MIXED if they possessed two regulated poly(A) sites that could either be located within the same exon or alternate final exons due to multiple isoforms, while genes with poly(A) sites located within internal exons or introns were termed internal APA (iAPA) (Figure S1B). Only in the case of TUTR genes can one be certain that APA results from the direct regulation of alternative poly(A) sites themselves, whereas all other events can result also from the regulation of alternative splice sites that are located between the poly(A) sites. Across these APA classes, the largest number of significant events (2779) were triggered by double ADMA and SDMA inhibition (Figure 1A, Figure S1C, supplemental Table 1). These events generally changed in the same direction also with individual inhibitors, with ADMA inhibition globally having a stronger impact than SDMA inhibition (Figure 1A, Figure 1B).

**Figure 1.**
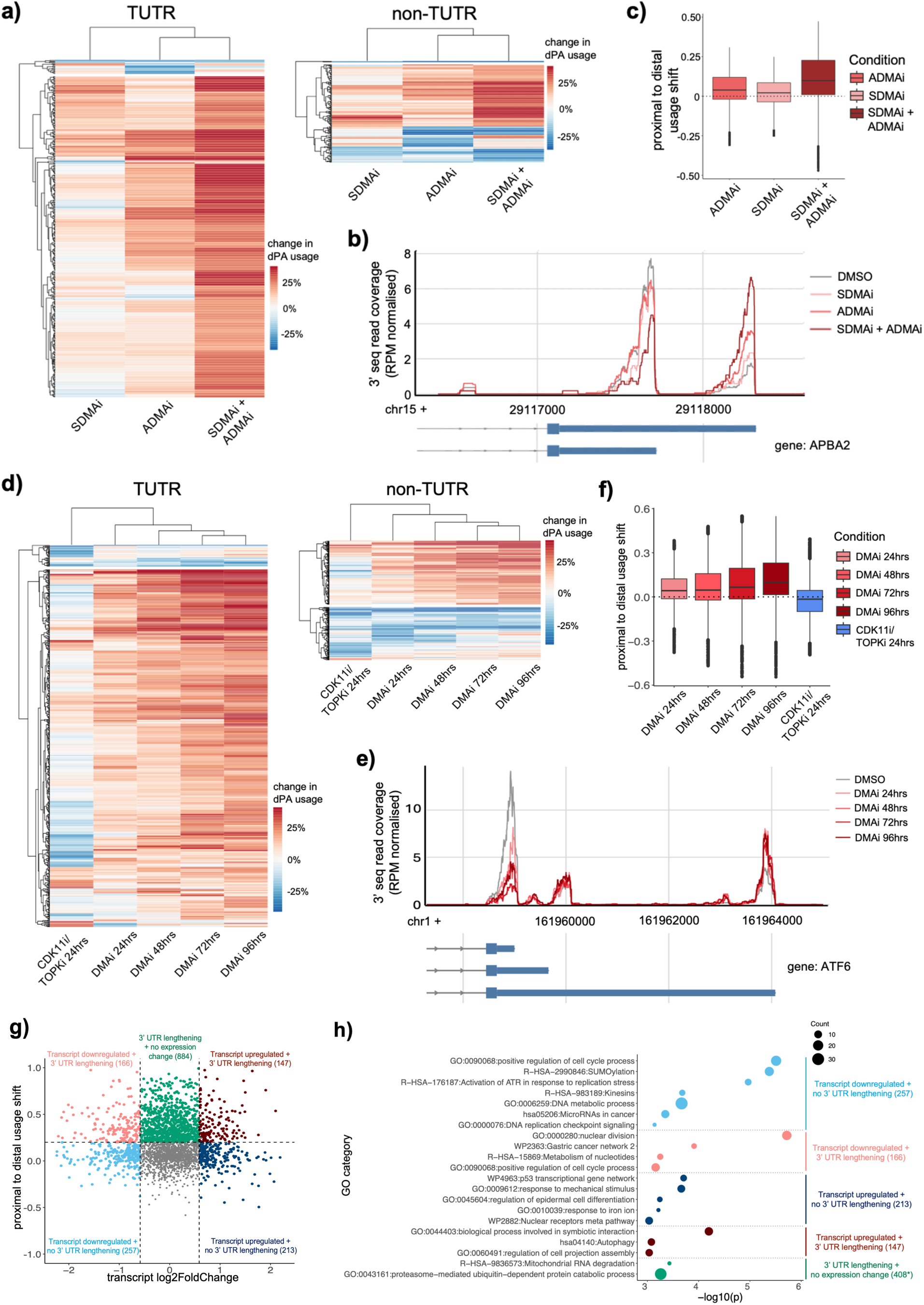
Inhibition of arginine methylation triggers 3’ UTR lengthening. **a)** Heatmaps displaying distal poly(A) site change in usage (%) upon SDMAi, ADMAi, or SDMAi+ADMAi relative to DMSO for significant TUTR (left) and non-TUTR (right) genes. **b)** An example of a gene displaying a significant proximal to distal poly(A) site switch following inhibition of arginine methylation. **c)** Distribution of proximal to distal poly(A) site usage shifts across all genes induced by DMAi treatment conditions. Proximal to distal usage shift is calculated as: dPA change in usage - pPA change in usage. **d)** Heatmap displaying distal poly(A) site change in usage (%) relative to DMSO for significant TUTR (left) and non-TUTR (right) genes. **e)** An example of a gene displaying a significant proximal to distal poly(A) site switch from 24 hours post-DMAi onwards. **f)** Distribution of proximal to distal poly(A) site usage shifts across all genes at each time point, calculated as in c). **g)** Scatter plot displaying changes in APA and gene expression following 96hrs SDMAi+ADMAi. **h)** Gene ontology term enrichment analysis of genes categorised according to their mode of regulation in response to arginine methylation inhibition. Selected set of significant terms is shown, with full set available in supplemental Table 3. * = proximal to distal usage shift threshold was increased to > 35% for the “3’ UTR and no expression change” group to ensure its size was more comparable to the other groups prior to the GO analysis.

A global shift toward proximal poly(A) (pPA) sites has been found in most cancer types, however the signalling pathways that may drive this shift remain unclear^33^. Strikingly, we found that arginine methylation strongly promotes such a shift. The vast majority of the significant APA events triggered by PRMT inhibition (ADMAi = 76.4%, SDMAi = 75.0%, SDMAi+ADMAi

= 93.5%) increased the relative use of the distal poly(A) (dPA) site (Figure 1A, Figure 1B). Indeed, expanding our analysis to all genes revealed a global trend of pPA site downregulation upon arginine methylation inhibition, particularly in the double SDMAi + ADMAi condition (Figure 1C). TUTR was the predominant significant APA event class identified in ADMAi and SDMAi+ADMAi conditions (Figure S1D), and had the most pronounced trend towards use of the dPA site (Figure S1E). However, in SDMAi-only conditions the proportion of splicing-related APA events (ALE category) was greater (Figure S1D), consistent with the previously described role of SDMA in spliceosome assembly and hence splicing regulation^19^. Thus, it is plausible that cellular signalling in cancer and other highly proliferating cells could act via arginine methylation to promote shorter 3’UTRs.

### 3’ UTR lengthening is a primary consequence of DMAi

To assess whether this 3’ UTR lengthening is likely a direct consequence of decreased arginine methylation, we assessed how quickly this occurs upon DMAi treatment. Given the strong increase in APA generated by combining inhibitors over and above single inhibition (Figure 1C), we chose double SDMA+ADMA inhibition (which we term ‘DMAi’) for this and all later experiments. We generated 3’ RNA-seq libraries from LU-99 cells at 24 hour intervals post-DMAi, starting at 24 hours through to 96 hours (Figure S2A). We observed clear APA shifts towards global 3’ UTR lengthening from the earliest DMAi time point tested (Figure 1D, Figure 1E, supplemental Table 2). Over 1000 significant events were identified already at the 24 hour time point (Figure S2B), and 77.2% increased the usage of dPA sites. Again, TUTRs were the most common class of significant events (Figure S2C), and 94.2% of significant TUTR events led to dPA upregulation (Figure S2D). The extent of proximal to distal poly(A) shift continued to increase at each later time point (Figure 1D, 1F), coinciding with the continued reduction in protein arginine methylation (Figure S2A).

In light of previous studies revealing that longer 3’ UTRs are associated with reduced proliferative states^28,29^, as well as literature demonstrating the efficacy of SDMA and ADMA inhibitors at slowing cancer cell growth^3,11^, we also assessed the possibility that the 3’ UTR lengthening observed following DMAi was a secondary consequence of reduced proliferation. For this purpose, we assessed APA in LU-99 cells treated with 50 nM TOPK and CDK11 inhibitor OTS964 for 24 hours, which reduces proliferation and downregulates cell cycle-related genes more strongly than DMAi at the equivalent time point (Figure S2E, S2F). We did not observe a global switch towards longer 3’ UTRs under this treatment (Figure 1D, Figure 1F). Instead, a slight increase in pPA site usage was observed. These findings indicate that reduced proliferation is unlikely the primary cause of the 3’ UTR lengthening in our cells.

### Functional implications of 3’ UTR lengthening

Ontology analyses of the genes containing the APA events significantly altered upon 24 hour DMAi identified several signalling and metabolic pathways that are often dysregulated within tumours (Figure S3A). For example, 24 hours DMAi downregulates the short mRNA isoform encoding the signalling factor FGF2 (Figure S3B), a potent modulator of proliferation; this short FGF2 mRNA isoform is recurrently induced across multiple cancers^34^. Similarly, 24 hours DMAi also downregulates the short mRNA isoform encoding the critical autophagy factor ATG7 (Figure S3C), with this proximal to distal poly(A) site switch reported to reduce tumour growth and be associated with significantly prolonged patient survival^35^. Thus, it is likely that 3’ UTR lengthening of mRNAs encoding proteins such as FGF2 and ATG7 contributes to the reduced proliferation of cells treated with DMAi.

We next studied the relationship between differential transcript abundance and TUTR-APA triggered by arginine methylation inhibition, focusing on 96 hours inhibition that exhibited the strongest phenotype. We did not observe any global correlation between the altered APA and transcript abundance, though a large number of genes were affected at both levels (Figure 1G). To classify the genes according to these changes, we set a threshold of 1.5-fold change for transcript abundance, and a proximal to distal usage shift > 20% for 3’ UTR lengthening. Ontology analyses revealed distinctions between the most enriched gene functions among these regulatory modes (Figure 1H). Genes with exclusively decreased transcript abundance, as well as those with 3’ UTR lengthening in addition to decreased transcript abundance, included primarily cell-cycle promoting ontologies (Figure 1H). Genes with exclusively increased transcript abundance included p53-network ontologies, genes with increased transcript abundance as well as 3’ UTR lengthening included autophagy-related ontologies, whereas genes with 3’ UTR lengthening but without altered transcript abundance included ubiquitin-related degradation ontologies (Figure 1H). Taken together, these results agree with recent findings that the regulation of APA and transcript-level abundance are generally independent^32,33^, though it is clear that the two mechanisms complement each other to implement the cellular effects of arginine methylation.

### DMA promotes short 3’ UTRs in highly proliferating cell types

Following the observation that DMAi induces 3’ UTR lengthening in mRNAs encoding several cancer- and proliferation-related proteins, we sought to establish whether this was specific to our MTAP-null lung cancer line, or whether it can be reproduced across cell lines with various genetics and tissue origins. Therefore we repeated the DMAi experiments on a panel of ten cancer lines across six tissues with varying p53 and MTAP status. Each line was subjected to 72 hour DMAi, after which we generated 3’ RNA-seq libraries. Global lengthening of 3’ UTRs was induced by DMAi in all cancer lines assayed (Figure 2A), with the trend especially apparent in TUTR genes (Figure 2B, Figure S4A). Moreover, most DMAi-triggered APA changes were reproducible across cell lines (Figure S4B), with 100s of poly(A) sites displaying significant change in usage in multiple cell lines (Figure 2B, Figure 2C). Indeed, we identified 768 ‘common DMAi’ poly(A) sites whose usage was altered by DMAi in every cell line in which they were present in the poly(A) site atlas (supplemental Table 4). These findings suggest that arginine methylation is required for the production of short 3’ UTR isoforms across cancer types.

**Figure 2.**
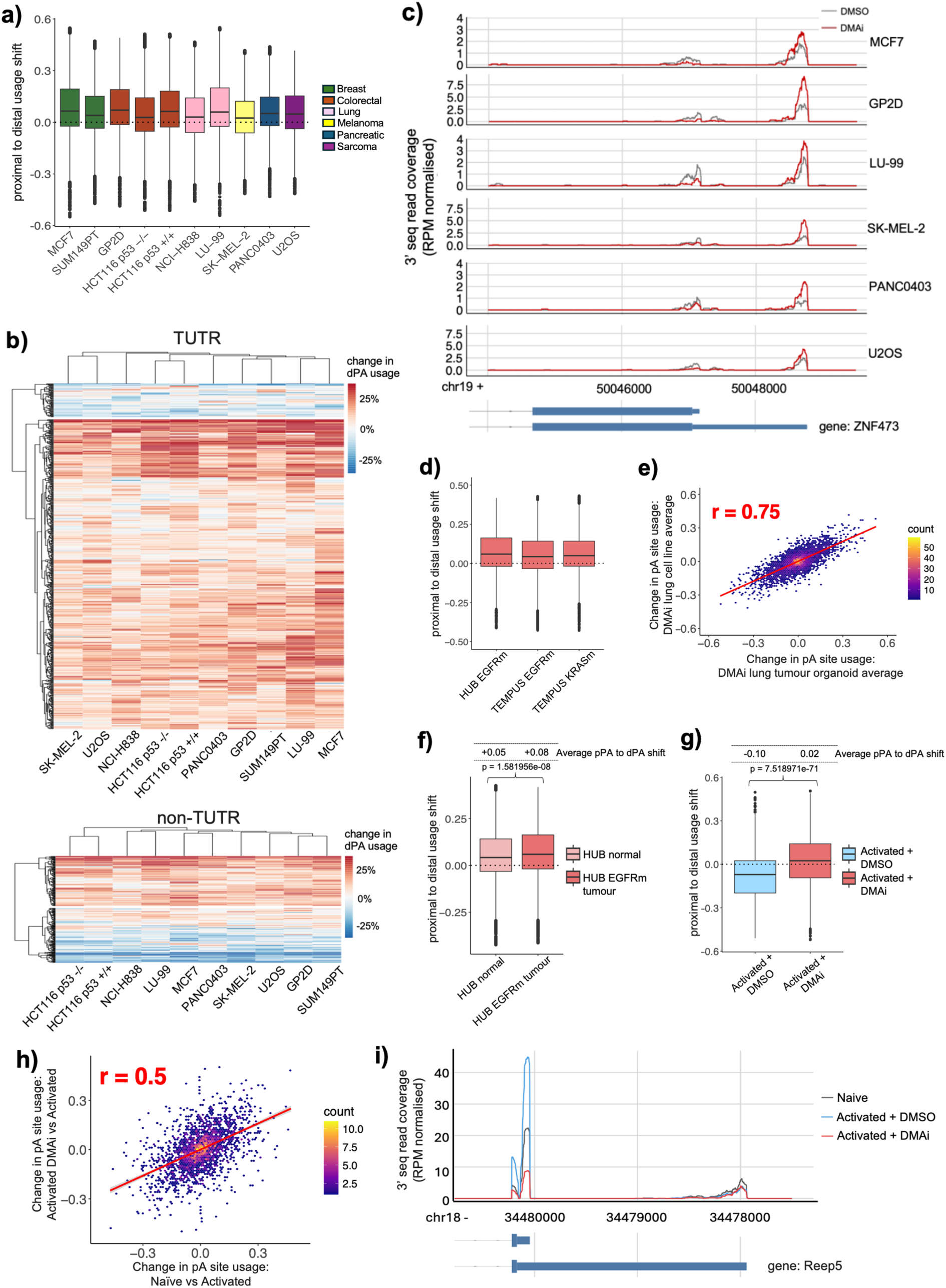
DMA maintains pPA site usage across multiple proliferating cell types. **a)** Distribution of proximal to distal usage shifts among all genes triggered by DMAi in each cancer line. Proximal to distal usage shift is calculated as: dPA change in usage - pPA change in usage. **b)** Heatmaps displaying DMAi-triggered distal poly(A) site change in usage (%) in significant TUTR (upper) and non-TUTR (lower) genes in each cancer line. Genes displayed in heatmap = genes for which change in usage data could be calculated in every line. **c)** An example of a DMAi-induced APA event conserved across multiple cell lines. **d)** Distribution of proximal to distal usage shifts among all genes triggered by DMAi across three lung tumour-derived organoids - HUB-07-B2-051 (EGFR mutant), TEMPUS AZ-574812 (EGFR mutant), TEMPUS AZ-291290 (KRAS mutant), calculated as in a). **e)** Scatter plot displaying Pearson correlation (r) in DMAi-induced poly(A) site usage change between patient-derived lung tumour organoids (averaged) and lung cancer lines (averaged). **f)** Distribution of proximal to distal usage shift among all genes triggered by DMAi in HUB-07-A2-051 normal lung tissue organoids (pink), or by DMAi in patient-matched HUB-07-B2-051 lung tumour-derived organoids (red), calculated as in a). Statistical significance was calculated using the Kruskal-Wallis Test. **g)** Distribution of proximal to distal usage shift among all genes in activated murine T cells (blue), or activated murine T cells also treated with DMAi (red), calculated as in a). Statistical significance was calculated using the Kruskal-Wallis Test. **h)** Scatter plot displaying Pearson correlation (r) of poly(A) site usage change between Naive vs Activated T cells and DMAi-treated Activated vs Activated T cells. **i)** An example of a 3’ UTR (*Reep5)* whose shortening upon T cell activation (blue) is prevented by DMAi (red).

We next examined the impact of DMAi on APA in four patient-derived lung organoid models - TEMPUS AZ-574812 (tumour, EGFR mutant), TEMPUS AZ-291290 (tumour, KRAS mutant), HUB-07-B2-051 (tumour, EGFR mutant) and HUB-07-A2-051 (derived from the tumour-proximal non-mutated cells of the same patient as HUB-07-B2-051). In accordance with our previous findings, DMAi triggered global 3’ UTR lengthening in all three lung tumour organoid models tested (Figure 2D), with the average DMAi-APA changes across these organoids demonstrating excellent correlation with the average DMAi-induced APA changes previously observed in our lung cancer cell line models (LU-99 and NCI-H838) (Figure 2E). When comparing baseline APA profiles between these patient-matched normal versus tumour organoids, we observed increased pPA usage in the tumour relative to normal (Figure S4C), consistent with the widespread literature documenting higher production of short 3’ UTR isoforms in the more highly proliferating cells of tumours^28,29^. DMAi triggered a significantly smaller amount of 3’ UTR lengthening in HUB-07-A2-051 normal organoids compared to the patient-matched HUB-07-B2-051 tumour organoids (Figure 2F). Thus, we find that the DMAi-induced 3’ UTR lengthening can be replicated in models that more accurately represent the 3D physiological architecture of tumours.

Given the consistent impact of DMAi upon APA across cancer cells regardless of their tissue of origin, we next examined whether this phenomenon could also be replicated in a non-pathological, high proliferative cellular context. We therefore proceeded to studies of activated T cells, which undergo increased proliferation that is coupled to global shortening of 3’ UTRs, much like cancer cells^28,36^. As expected, we observed global 3’ UTR shortening when naïve murine CD4+ T cells were activated by treatment with anti-CD3, anti-CD28 and rIL-2, but this was completely abolished if DMAi was included during T cell activation (Figure 2G). Globally, the APA changes of naive versus activated T cells positively correlated with the APA changes of DMAi+activated versus activated T cells (Figure 2H). Therefore, DMAi strongly blocks the changes in APA and thus prevents the 3’ UTR shortening of transcripts during T cell activation. For example, DMAi treatment fully blocks the shortening of genes such as *Reep5* and *Prkca* (Figure 2I, Figure S4D), which play critical roles in endoplasmic reticulum organization and signal transduction respectively, and have previously been shown to undergo 3’ UTR shortening upon T cell activation^36^.

In summary, our data indicate that protein arginine methylation is required for the 3’ UTR shortening that accompanies increased proliferation across a broad range of cell types.

### DMAi mitigates 3’ UTR shortening induced by depletion of CFIM

Cleavage and polyadenylation is a co-transcriptional process, and since proximal poly(A) sites are the first to be transcribed, they have the temporal advantage for CPA assembly. However, their core sequence motifs such as the canonical polyadenylation signal AAUAAA, upstream UGUA motifs, and downstream GU-rich sequences, which are all recognised and bound by CPA complexes, generally have weaker representation and greater deviation from the consensus compared to distal poly(A) sites. When functionality of the CPA machinery is compromised either by chemical inhibition or reduced expression of its core components, usage of distal poly(A) sites is generally increased, presumably because the sequence advantage of the distal sites becomes a more important factor than the temporal advantage of the proximal sites^37–41^. An exception is depletion of CFIM25 (also NUDT21) or CPSF6 (also CFIM68), two components of the cleavage factor I complex (CFIM) that specifically recognises the UGUA motifs. Reduced expression of CFIM25 was found to result in widespread 3’ UTR shortening due to the decreased capacity of the CPA machinery for recognising UGUA-rich distal sites^42,43^. These shortened isoforms are in many cases oncogenic, as they result in elevated expression of pro-proliferative factors^43–46^.

We hypothesized that DMAi might mitigate the oncogenic 3’ UTR shortening by counteracting the impact of the CFIM loss. Therefore we next tested how DMAi treatment alters the APA changes induced by an siRNA-mediated depletion of CFIM25 in LU-99 cells. In agreement with previously published findings^42,43^, we observed widespread downregulation of distal poly(A) sites upon 72 hour treatment with siRNA against CFIM25 (siCFIM25) (Figure 3A) and enrichment of UGUA motifs upstream of these sites (Figure S5A). However, this 3’ UTR shortening was mitigated if we included DMA inhibitors during the 72 hour siCFIM25 period (Figure 3A). The extent of mitigation differed between transcripts; therefore we grouped transcripts according to the level of DMAi-induced mitigation: full mitigation (189 transcripts), partial mitigation (384 transcripts) and no mitigation (235 transcripts) (Figure 3B), and conducted a motif enrichment analysis. This revealed an enrichment of GC-rich motifs in the full-mitigation transcripts at both proximal and distal poly(A) sites (Figure S5B). Indeed, the incidence of GC-rich motifs was strongly linked with the degree of DMAi mitigation, with the full-mitigation transcripts displaying elevated frequency of these motifs over several hundred nucleotides around the regulated poly(A) sites (Figure S5C). In light of this finding, we assessed GC-content across the entire gene region for each transcript group. This revealed that the increased GC-density of DMAi-sensitive transcripts is a gene-wide phenomenon, and thus might involve broader transcriptional and chromatin mechanisms (Figure 3C).

**Figure 3.**
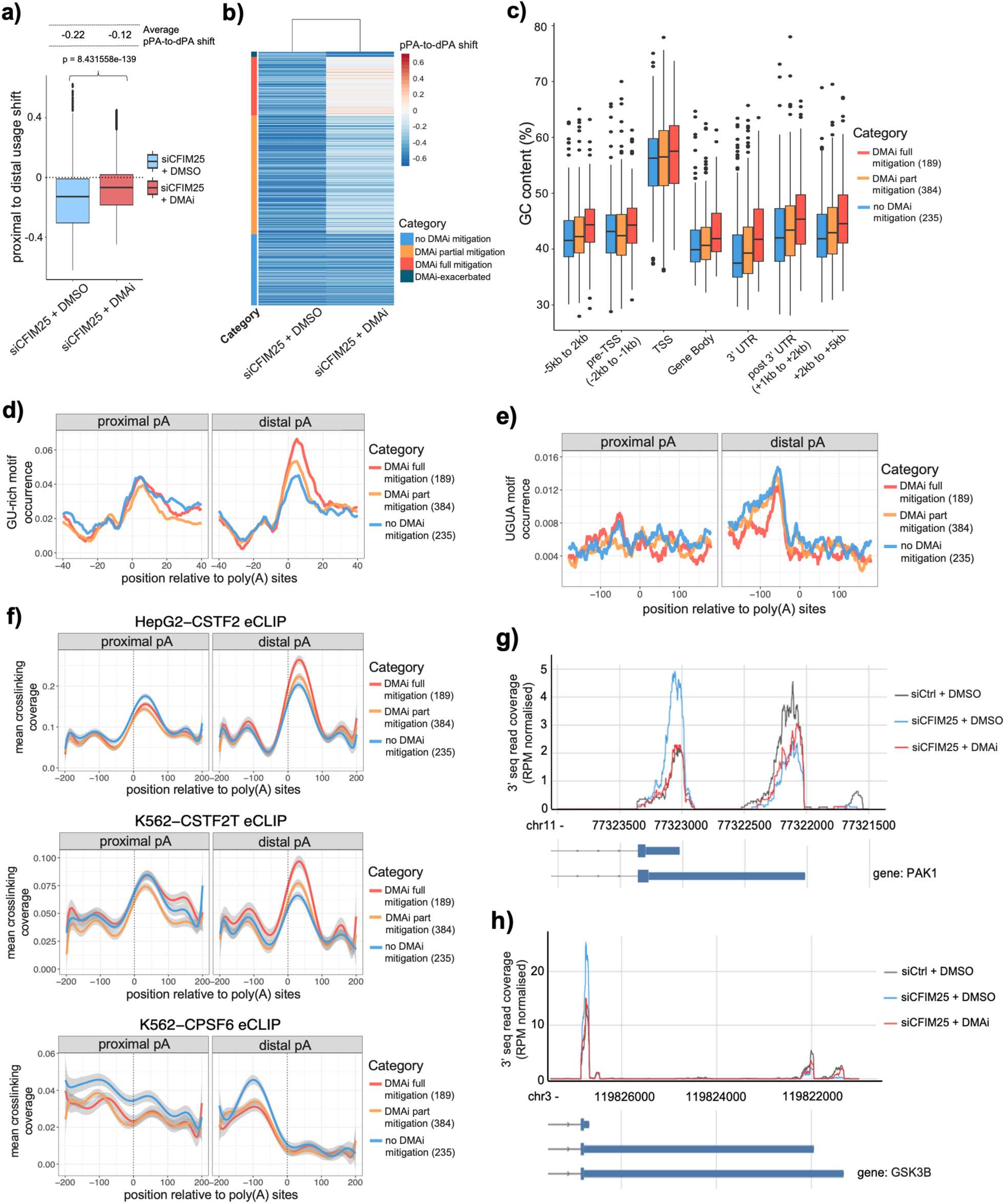
3’ UTR shortening induced by CFIM25 loss is alleviated by DMAi. **a)** Distribution of proximal to distal poly(A) site usage shifts across all genes triggered by siCFIM25 (blue), or siCFIM25 in combination with DMAi (red). Proximal to distal usage shift is calculated as: dPA change in usage - pPA change in usage. Statistical significance was calculated using the Kruskal-Wallis Test. **b)** Heatmap showing proximal to distal poly(A) usage shifts in siCFIM25-shortened 3’ UTRs, categorised according to the degree of mitigation achieved by DMAi. Full DMAi mitigation: siCFIM25 + DMAi prox-to-dist shift <20% of siCFIM25 + DMSO prox-to-dist shift. Partial DMAi mitigation: siCFIM25 + DMAi prox-to-dist shift between 30%-60% of siCFIM25 + DMSO prox-to-dist shift. No DMAi mitigation: siCFIM25 + DMAi prox-to-dist shift >80% of siCFIM25 + DMSO prox-to-dist shift. DMAi exacerbated: siCFIM25 + DMAi prox-to-dist shift >133% of siCFIM25 + DMSO prox-to-dist shift. **c)** Distribution of % GC content within each gene region among each DMAi sensitivity category. **d)** Comparison of position-dependent frequency of occurrence of GU-rich motifs in the regions centred on pPA or dPA at DMAi-full mitigation (red), partial mitigation (orange) and no mitigation (blue) siCFIM25-shortened 3’ UTRs. Running averages were calculated over 15 consecutive nucleotide positions. GU-rich motif = any 4mer combination of 2Gs + 2Us **e)** Comparison of position-dependent frequency of occurrence of UGUA within proximal and distal regions in 3’ UTRs across each DMAi sensitivity category. Running averages were calculated over 30 consecutive nucleotide positions. **f)** Upper = CSTF2 eCLIP binding around poly(A) sites in 3’ UTRs across each DMAi sensitivity category (expression matched), calculated from ENCODE consortium data in HepG2. Middle = CSTF2T eCLIP binding around poly(A) sites in 3’ UTRs across each DMAi sensitivity category (expression matched), calculated from ENCODE consortium data in K562. Lower = CPSF6 eCLIP binding around poly(A) sites in 3’ UTRs across each DMAi sensitivity category (expression matched), calculated from ENCODE consortium data in K562 cells. **g)** Visualisation of Pak1 3’ UTR proximal and distal poly(A) site usage upon siCFIM25 and siCFIM25 + DMAi. **h)** Visualisation of GSK-3β 3’ UTR proximal and distal poly(A) site usage upon siCFIM25 and siCFIM25 + DMAi.

We then analysed the composition of the core CPA binding motifs across transcript groups. While the strength of poly (A) signal upstream of poly(A) sites was found to be very similar across all transcript groups (Figure S5D), DMAi-full mitigation transcripts displayed highly elevated levels of GU-rich motifs immediately downstream of distal poly(A) sites (Figure 3D), highlighting strong distal Downstream Sequence Elements (DSEs) as an additional characteristic of DMAi-sensitive genes. In contrast, the frequency of upstream UGUA motifs inversely correlated with DMAi-sensitivity (Figure 3E). Thus, while DMAi cannot rescue the distal sites that most strongly depend on the recognition of UGUA motifs by CFIM25, it can at least partially rescue most other distal sites.

To identify factors that might mediate the DMAi effects, we compared the crosslinking profiles determined by the eCLIP experiments of 150 RBPs that are available from the ENCODE project around the proximal and distal poly(A) sites within each transcript group. The strongest peak downstream of poly(A) sites was observed for CSTF2 (also known as CSTF64) and CSTF2T eCLIP experiments (Figure S5E), which overlapped with the expected position of DSEs. Notably, the CSTF2 and CSTF2T binding was strongest downstream of the distal poly(A) sites of full-mitigation transcripts (Figure 3F, Figure S5F). Conversely, several eCLIP datasets of other RBPs had reduced crosslinking profiles around the poly(A) sites of full-mitigation transcripts, including CPSF6 and KHSRP (Figure 3F, Figure S5G). This could not be simply a consequence of pPA/dPA usage ratios, as the full-mitigation transcripts on average had intermediate ratios between partial- and no-mitigation groups (Figure S5H), whereas the partial-mitigation group had intermediate RBP binding profiles. Thus, DMAi is most efficient in promoting lengthening of 3’ UTRs where the recognition of distal poly(A) sites depends more on the CSTF than the CFIM complex. This is evident by the high incidence of the CSTF-binding GU-rich motifs and the high eCLIP signal for CSTF components, and by the relatively low incidence of CFIM-binding UGUA motifs and the low eCLIP signal for CPSF6. Notably, previous studies have demonstrated that CSTF components are direct targets of arginine dimethylation^3,25,47^. Our results indicate that arginine methylation of CSTF complex components decreases the usage of distal GU-rich poly(A) sites, and this is required for the global 3’ UTR shortening even if the activity of the CFIM complex is decreased.

We found that the DMAi fully-mitigated transcripts encode numerous proteins with important functions in cancer biology. This includes Pak1, a key component of the Ras signalling pathway and GSK-3β, a serine/threonine protein kinase that has emerged as a promising cancer target^45,48^. DMAi totally prevents the 3’ UTR shortening of Pak1 and GSK-3β mRNAs upon CFIM25 depletion (Figure 3G, 3H), which was reported in both cases to result in higher protein expression, and in the case of Pak1 also associated with poor patient prognosis^45^. Thus, targeting protein arginine methylation enables the complete prevention of many oncogenic consequences triggered by reduced CFIM25 expression, itself a phenomenon reported across many tumour types.

### DMAi-induced APA shares transcriptomic signatures with CPA disruption

To further understand the mechanisms that characterise DMAi-induced APA, we compared this phenotype with APA triggered by other cellular perturbations. Knockdown or inhibition of core CPA factors can induce global changes in APA, with two such perturbations - CPSF73 inhibition (CPSF73i) and PCF11 knockdown (siPCF11) - inducing especially widespread 3′ UTR lengthening in cancer cells and have hence been suggested as potential therapeutic strategies^41,49,50^. In addition to these primary APA factors, auxiliary RBPs such as KHDRBS1^51^ and ELAVL1^52^ can also modulate APA at specific transcripts, often by enhancing or antagonizing the core CPA machinery. Therefore, we collated a panel of previously published or internally generated 3’ RNA-seq datasets comprised of these various APA-inducing cellular perturbations, including CPSF73i and siPCF11 (as well as several other RBP knockdowns, enzymatic inhibitions etc.), and analysed them via the same APA/DRIMSeq pipeline that we have applied to the study of DMAi. Among these datasets, those that triggered 3’ UTR lengthening by directly disrupting the CPA machinery - CPSF73 inhibition and PCF11 knockdown - showed strong positive correlation with DMAi-induced APA (Figure 4A), with siPCF11 exhibiting the highest correlation overall (Figure 4A). Given that PCF11 is a known arginine-methylation substrate^53^, this finding supports the notion that DMAi triggers APA principally through its direct impact on CPA protein function.

**Figure 4.**
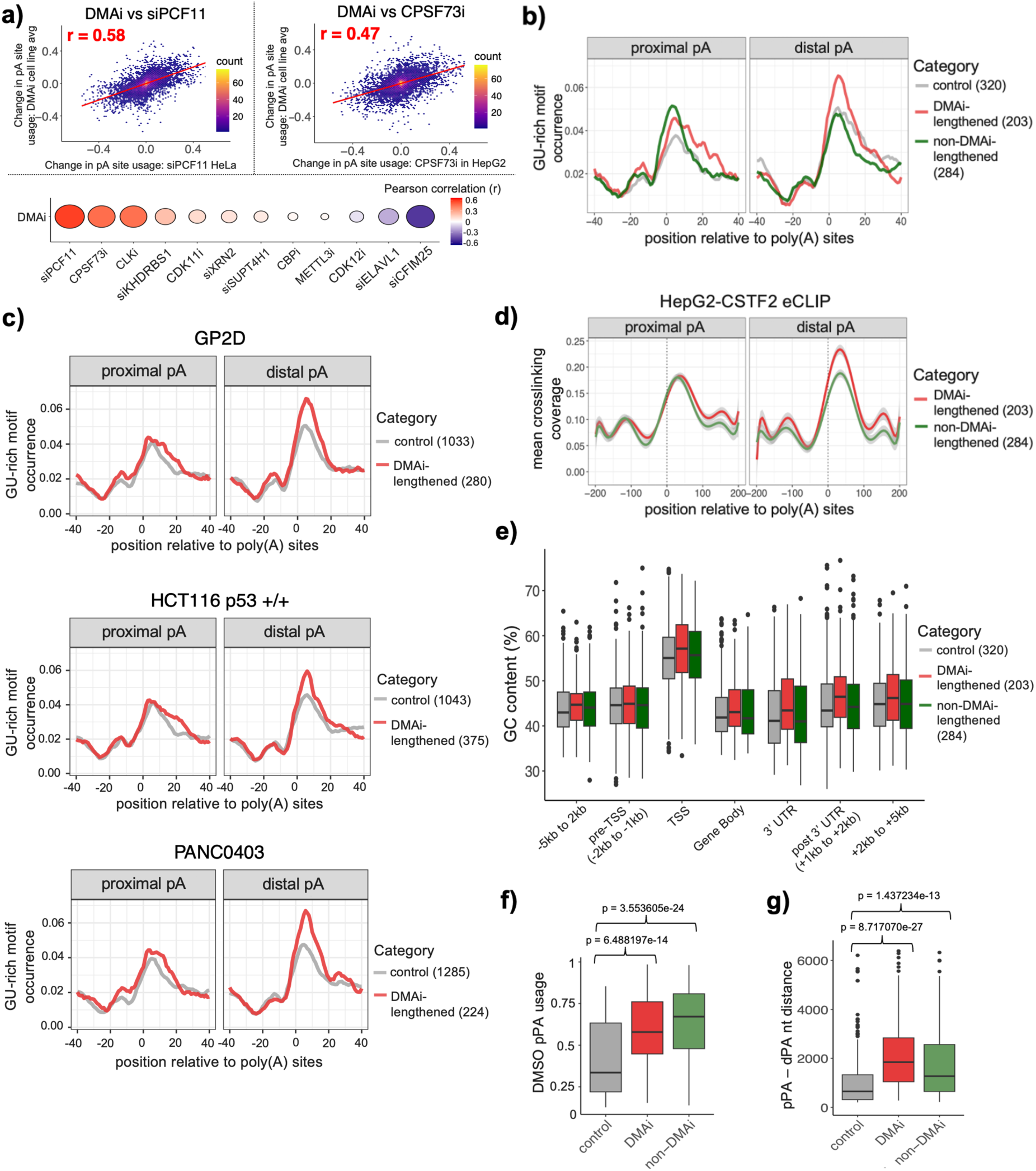
DMA-regulated 3’ UTRs possess strong distal DSEs and are GC-rich. a) Upper = Scatter plots displaying Pearson correlation (r) in poly(A) site usage change triggered by DMAi versus siPCF11 (left) and CPSF73i (right). Lower = Pearson correlation (r) in poly(A) site usage change triggered by DMAi versus all other APA-inducing perturbation datasets within the 3’ RNA-seq panel. **b)** Comparison of position-dependent frequency of occurrence of GU-rich motifs within proximal and distal regions at DMAi-lengthened (red), non-DMAi-lengthened (green) and control (grey) sites. Running averages were calculated over 15 consecutive nucleotide positions. GU-rich motif = any 4mer combination of 2Gs + 2Us. **c)** Comparison of position-dependent frequency of occurrence of GU-rich motifs within proximal and distal regions at DMAi-lengthened (red) and unchanged, control (grey) 3’ UTRs, across three representative cancer lines. Upper = GP2D, Middle = HCT116 p53 +/+, Lower = PANC0403. **d)** CSTF2 eCLIP binding around poly(A) sites in 3’ UTRs across DMAi-vs non-DMAi-lengthened transcripts (expression matched), calculated from ENCODE consortium data in HepG2 cells. **e)** Distribution of % GC content within each gene region among DMAi-lengthened (red), non-DMAi-lengthened (green) and control (grey) genes. **f)** Distribution of baseline proximal poly(A) site usage in control (grey), DMAi-lengthened (red) and non-DMAi-lengthened (green) categories (expression matched). Scale: 0 = 0% usage, 1 = 100% usage. Statistical significance was calculated using the Kruskal-Wallis Test, with Bonferroni method p value adjustment for multiple testing. **g)** Boxplot showing the nucleotide distance between proximal and distal poly(A) sites at control (grey), DMAi-lengthened (red) and non-DMAi-lengthened (green) sites. Statistical significance was calculated using the Kruskal-Wallis Test, with Bonferroni method p value adjustment for multiple testing.

Across our entire panel of 3’ RNA-seq datasets we identified hundreds of transcripts that are 3’ UTR lengthened by at least one non-DMAi-related perturbation, but not by DMAi; which we refer to as non-DMAi transcripts. To further dissect the features that confer 3′ UTR sensitivity to DMAi, we proceeded to compare these non-DMAi transcripts with our previously identified ‘common DMAi’ transcripts that were lengthened by DMAi treatment across multiple cancer lines. Motif enrichment analyses comparing these two transcript groups identified GU-rich motifs as especially prevalent at the DSE of distal poly(A) sites in DMAi-lengthened transcripts (Figure S6A), and plotting their incidence around these regions confirmed their enrichment in DMAi vs non-DMAi transcripts (Figure 4B). This enrichment was also apparent when comparing DMAi-lengthened versus unchanged control transcripts within individual cancer cell lines (Figure 4C). This again is reflected by elevated CSTF2 eCLIP signal at DMAi-regulated DSEs compared to non-DMAi sites (Figure 4D), while other RBPs do not show any clear signal differences (Figure S6B).

As previously, the frequency of canonical poly(A) signals at distal poly(A) sites did not differentiate DMAi-regulated vs non-DMAi-regulated transcripts (Figure S6C). We next analysed GC-content, which once again revealed increased GC-abundance in DMAi-sensitive compared to controls and non-DMAi-sensitive transcripts (Figure 4E). In line with earlier findings, this increased GC-content spanned across the whole genes (Figure 4E), and the same enrichment was obtained if DMAi-sensitive versus control transcripts were compared in each cancer line individually (Figure S6D). High GC-content has previously been reported as a molecular signature of genes that display increased sensitivity to compromised CPA machinery^41^, and indeed we observe this GC-rich bias within transcripts that are lengthened both in the CPSF73i and the siPCF11 datasets (Figure S6E, S6F). These commonalities between the impact of DMAi or perturbed CPA machinery on APA suggest that arginine methylation might act by increasing the efficiency of the CPA machinery.

We also identified additional features that were characteristic both of the DMAi-lengthened and non-DMAi lengthened 3’ UTRs, including high baseline proximal poly(A) site usage (Figure 4F), as well as greater distance between proximal and distal poly(A) sites compared to their unchanged counterparts (Figure 4G). In light of this finding, we assessed 3’ UTR length in our previously identified siCFIM25-shortened transcripts categorised by DMAi-sensitivity, and again observed that the DMAi full-mitigation transcript group had a greater distance between poly(A) sites compared to the remaining siCFIM25-shortened transcripts (Figure S6G). In summary, these findings demonstrate that the transcripts that undergo DMAi-induced 3’ UTR lengthening contain key hallmarks, many of which are shared with other CPA-perturbing strategies. This informs on the likely direct impact of arginine methylation on the 3′ end processing machinery, and suggest a potential for its synergy with other APA-directed therapeutic interventions.

## DISCUSSION

Our findings reveal protein arginine methylation as a critical modulator of APA, maintaining shortened 3’ UTR isoforms across a broad range of proliferating cell types. Single inhibition analyses demonstrate that ADMA has a stronger regulatory effect than SDMA, suggesting that RBP substrates of PRMT1 rather than PRMT5 are the primary mechanistic actors driving this pathway. However, the strong increase in 3’ UTR lengthening phenotype triggered by double inhibition indicates that SDMA can strongly compensate in the absence of ADMA, either by methylating the same substrates through substrate scavenging, or by methylating other proteins that become more essential for pPA recognition in the absence of ADMA. This pronounced 3’ UTR lengthening upon double DMAi could help explain the synergistic effects reported when treating cancer cells with both inhibitors^11^.

APA transcripts lengthened by DMAi or by compromised CPA activity shared many features, indicating that arginine methylation promotes high efficiency of the CPA machinery required for recognition of proximal poly(A) sites at these genes. This is exemplified by the strong correlation observed between DMAi-induced APA and APA induced by knockdown of PCF11 - an ADMA-methylated CFIIM component - whose substrate arginines are located within FEGP repeats necessary for the RNA cleavage activity of this complex in vitro^54^. Ubiquitinylation of PCF11 has also been shown to regulate the interactions between RNA polymerase II and other components of the CPA machinery and hence APA^55^, further highlighting the impact of post-translational modifications upon the assembly of the multi-complex CPA apparatus.

Both DMAi and compromised CPA activity affect transcripts with long 3’ UTRs in GC-rich genes that usually favour high proximal poly(A) site usage in proliferating cells. We found that GC-richness is a feature of these genes as a whole, rather than being restricted to the regions around the regulated poly(A) sites, which indicates that it may impact the chromatin state and transcription of these genes, and therefore the co-transcriptional nature of CPA regulation is likely to be important. Moreover, the observation that the two tandem poly(A) sites are further apart than in unaffected 3’ UTRs indicates that the temporal advantage of the proximal sites being transcribed first is important for these sites, because the spacing between the two sites affects the extent of the co-transcriptional advantage.

An additional hallmark of DMAi-lengthened mRNAs is strong GU-rich distal DSEs, which facilitate high affinity, stable interactions with components of the CSTF complex^37^. Since CSTF2 is arginine methylated^3,25,47^, this further suggests that arginine methylation changes the co-transcriptional activity of the CPA machinery such that proximal poly(A) sites can be processed despite the sequence superiority of these distal poly(A) sites. While the CSTF complex is mechanistically implicated by this finding, DMAi-induced 3′ UTR lengthening encompasses a broader class of transcripts than CSTF2/CSTF2T knockdown alone^37^, consistent with the fact that multiple other core CPA components are also arginine methylated. It is thus likely that the full mechanism of arginine methylation on APA involves a cumulative interplay of many modified components and their interactions with each other and other machineries, especially the RNA polymerase II that is also ADMA-modified at its C-terminus^56^.

Recent years have witnessed a growing appreciation for altered APA as a driver of tumourigenesis in many cancers. Mounting evidence shows that cancer-linked APA events strongly correlate with altered expression of core APA factors - changes that not only trigger oncogenic 3′ UTR shortening but also serve as accurate predictors of patient prognosis and tumour aggressiveness^45,57^. For example, downregulation of CFIM25 in various tumours leads to widespread 3′ UTR shortening (including in the PAK1 transcript)^43,45^, whereas CSTF2 upregulation likewise accelerates the production of shorter, more oncogenic mRNA isoforms^58^. Our results clearly demonstrate that PRMT inhibition can reverse numerous oncogenic APA isoforms, including those triggered by reduced expression of CFIM25. Therefore, future studies could assess the potential for synergistic activities of PRMT inhibitors with other pharmacological modulators of APA, such as JTE607-mediated CPSF73 inhibition^50,59^. This strategy might help overcome the therapeutic window limitations seen in prior PRMT inhibitor trials^17,18^ and pave the way for more targeted, combination-based therapies that exploit the unique vulnerability of APA-addicted tumours.

Our observation that PRMT inhibition also blocks the shift to shorter 3′ UTRs in activated T cells underscores the fact that arginine methylation is crucial not only in pathologically proliferating tumour cells but also in physiological high-proliferation states. This highlights PRMT-mediated arginine methylation as a fundamental requirement for robust 3′ end processing across diverse cellular contexts. Moreover, it suggests that targeting PRMT activity could have therapeutic relevance not only for malignancies but also for diseases characterized by elevated T cell proliferation or dysregulated immune responses.

## ACKNOWLEDGEMENTS

The authors would like to thank the following Francis Crick Institute Science Technology Platforms for their technical support: Advanced Sequencing Facility, High-Performance Computing and Data Analytics Platform (NEMO), and Cell Services. This work was supported by the UK Dementia Research Institute [award number UK DRI-RE21605] through UK DRI Ltd, principally funded by the UK Medical Research Council; by the Francis Crick Institute which receives its core funding from Cancer Research UK (CC0102), the UK Medical Research Council (CC0102), and the Wellcome Trust (CC0102). LG was a recipient of post-doctoral funding by AstraZeneca. AMC is supported by a Wellcome Early Career Award (225081/Z/22/Z).

## COMPETING INTERESTS

SM, AF, JTL and JU are current or former employees and shareholders of AstraZeneca.

## METHODS

### Cell culture

HCT116 p53 +/+, HCT116 p53 -/- and MCF7 cells were obtained from Cell Services at The Francis Crick Institute and cultured in DMEM+GlutaMAX (Thermo Scientific) + 10% FBS. GP2D, NCI-H838, LU-99, PANC0403, SK-MEL-2, SUM149PT and U2OS cells were sourced from the AstraZeneca Global Cell Bank. GP2D, NCI-H838, LU-99 and PANC0403 cells were cultured in RPMI + 10 % FCS + 1 % Glutamine. SK-MEL-2 cells were cultured in EMEM + 10 % FCS + 1 % Glutamine. SUM149PT cells were cultured in Ham’s F-12 + 10 % FCS + 1 % glutamine. U2OS cells were cultured in McCoys + 10 % FCS + 1 % Glutamine. All cells were grown at 37 °C, 5 % CO2. For all arginine methylation inhibition experiments, cells were treated with 100 nM LLY-283 to inhibit SDMA and 3 μM GSK3368715 to inhibit ADMA.

### T cell isolation and activation

Spleens and axillary, brachial, mesenteric and inguinal lymph nodes of female C57/Bl6 mice were extracted, then homogenized through a 70 mm filter in RPMI 1640. Resulting cell pellets were then treated with 3ml ACK red blood cell lysis buffer. CD4+ T cells were then purified via MACS column (Miltenyi Naive CD4+ T Cell Isolation Kit, mouse, catalogue number 130-104-453) resulting in purity of >95%. 1,000,000 naïve CD4+ T cells were plated in flat bottom 96 well plates coated with 10 μg/ml anti-CD3 (clone 145-2C11), then treated with 4 μg/ml anti-CD28 (37.51) and 10 U/ml rIL-2 (Peprotech) in 200 μl total volume RPMI 1640. To inhibit arginine methylation, 100 nM LLY-283 and 3 μM GSK3368715 was added alongside anti-CD28 and rIL-2. Cells were harvested after 72 hours.

### Patient-derived organoids

The following organoid models were used within AstraZeneca: HUB-07-A2-051 (normal lung), HUB-07-B2-051 (matched tumour lung to normal organoid model 07-A2-051), TEMPUS AZ-574812 (tumour lung), TEMPUS AZ-291290 (tumour lung). HUB-07-A2-051 and HUB-07-B2-051 were procured from the Foundation Hubrecht Organoid Biobank under a license from HUB Organoids, while models AZ-574812 and AZ-291290 were acquired from Tempus Labs^60^. Organoids were grown according to supplier’s instructions (proprietary from HUB or TEMPUS, as per respective model) and passaged by mechanical dissociation into wells of 6 well plates (homogeneous populations below 100um). Organoids were allowed to settle in full media with RockI for 48 hours before compound was dosed in media without RockI. Models were treated with 100 nM LLY-283 and 3 μM GSK3368715 in combination for 48 hours (or a DMSO control) before harvesting. RNA extraction from at least 3 biological replicates was carried out using the RNeasy kit (Qiagen) according to manufacturer instructions and shipped to KCL for further processing.

### Cell viability assay

Cell viability was calculated through quantifying intracellular ATP concentrations using the CellTiter-Glo® 2.0 Cell Viability Assay, according to the manufacturer’s guidelines (Promega). Bioluminescence counts were quantitated using a CLARIOstar reader (BMG Labtech). Viability was expressed as a percentage of DMSO-treated cell viability.

### siRNA transfection

siRNA-induced knockdown was achieved using Silencer™ Select siRNAs (Thermofisher) combined with Lipofectamine™ RNAiMAX Transfection Reagent (Thermofisher) following the manufacturer’s forward transfection protocol. All siRNA transfections were carried out for 72 hours, at a final concentration of 10 nM. siRNAs used: Silencer™ Select Negative Control No. 1 siRNA (Thermofisher,), Silencer™ Select NUDT21 siRNA (Thermofisher, ID: s21772), Silencer™ Select ELAVL1 siRNA (Thermofisher, ID: s4608), Silencer™ Select KHDRBS1 siRNA (Thermofisher, ID: s20953). For analyses assessing the combined effects of siCFIM25 and DMAi, methylation inhibitors were added 24 hours after the siRNA, so that 48 hours inhibition had occurred at the point of harvesting.

### Western Blotting

Protein concentrations were calculated using the DC protein assay kit (BioRad). Lysates were supplemented with 4X NuPage loading buffer (+ 1mM DTT) and separated over 4–12% gradient SDS-PAGE gels. Proteins were then transferred to a 0.2 μm nitrocellulose membrane via the Trans-Blot Turbo RTA Mini Nitrocellulose Transfer Kit (Bio-Rad). The membranes were blotted with the primary antibodies: anti-SDMA (Cell signalling, #13222) at 1:1000, anti-ADMA (Cell signalling, #13522) at 1:1000, anti-actin at 1:5000 (abcam, ab8226). Blots were incubated with primary antibodies at 4°C overnight. The following day, LI-COR secondary Antibodies (IRdye680 1:15000, IRdye800 1:15000) were used to detect signals that were then visualized by Odyssey scanning (LI-COR).

### RNA extraction

Cells were washed once in 1x PBS and pelleted. RNA extractions were then performed on pellets using the Maxwell RSC simplyRNA kit (Promega) on a Maxwell RSC Instrument (Promega) according to the manufacturer’s instructions.

### 3’ RNA-seq

500 ng RNA was fragmented in 10 mM Tris-HCl pH 7.5, 3 mM MgCl2 buffer at 95°C for 12 minutes (final volume = 4 µL). 0.5 µL 5 µM oligo-dT reverse transcription primers containing Illumina P7 sequences and 0.5 µL 10 mM dNTPs were added to the fragmented RNA, before heating at 65°C for 3 min, then cooling to 42°C at 1°C/s. Reverse transcription was then performed using SuperScript IV (Invitrogen) according to manufacturer instructions, with the addition of 0.25 µL 40 µM template-switching oligo containing UMIs and Illumina P5 sequences. Following 1 hour incubation at 42°C, cDNA was purified using Mag-Bind TotalPure NGS beads. Libraries were generated using 0.5 µM i5/i7 Illumina indexed primers in Phusion High-Fidelity PCR Master Mix with HF buffer. Three replicates per condition were generated in all experiments, with the exception of the murine T cell studies for which four replicates per condition were generated.

### PolyA site usage analysis

The nf-quantseq pipeline (v.0.1.0; DOI: 10.5281/zenodo.14417140) was utilised to pre-process, deduplicate and map raw reads against the GRCh38 Homo sapiens (with the GENCODE version 45 annotation) or GRCm39 Mus musculus (with the GENCODE version M34 annotation) genome, as well as to generate a poly(A) site atlas containing all detectable sites across all same species experiments. BEDtools was employed to tally the number of reads mapping to a window 200 nucleotides upstream of each poly(A) site in a strand-aware manner. The count table generated served as input for the DRIMSeq package^32^ to identify differential poly(A) site usage between conditions. Pairwise comparisons were performed between DMSO and methylation inhibitor-treated cells in all experiments. Genes were required to have at least 10 reads in 75% of all samples, and poly(A) sites were required to have at least 5 reads in 75% of samples in the smallest group size, ensuring reliable detection across conditions. DRIMSeq was used to fit gene-level Dirichlet-multinomial models and feature-level beta-binomial models. A likelihood ratio test assessed differential poly(A) site usage between conditions. To control for multiple testing and maintain a gene-level false discovery rate of 5%, we employed a two-stage testing procedure using the stageR package^61^. Adjusted p-values were obtained, and poly(A) sites with adjusted p-values less than 0.05 and that exhibited an absolute change in usage of at least 10% between conditions were deemed significantly altered poly(A) sites. Metascape^62^ was utilised to identify enriched Gene Ontology terms and KEGG pathways among significantly altered poly(A) sites. 3’ RNA-seq read coverage at 3’ UTRs was visualised using clipplotr^63^.

### Differential gene expression analysis

The nf-core/rnaseq (v3.14.0) pipeline was used to pre-process and map raw reads against the GRCh38 Homo sapiens genome (with the GENCODE version 45 annotation), specifying no read length correction as recommended for 3’ RNA-seq data. Salmon quantification output files were imported into R using tximport. Differential expression analysis was then performed using DESeq2^64^, with effect size shrinkage using the apeglm package^64,65^. Metascape^62^ was utilised to identify enriched Gene Ontology terms and KEGG pathways among significant genes.

### Classification of APA events

The two poly(A) sites with the highest usage per gene were selected for the APA event categorisation, or in the case of genes with multiple significantly changed poly(A) sites, the two sites with the largest change in usage were selected. These two poly(A) sites for each gene were then assigned to transcript isoforms based on proximity between the polyA site coordinate and the end of the transcript, as defined in the gencode.v45.annotation.gtf annotation file. Transcript isoforms were interrogated for the presence of a splice site located between the proximal poly(A) site and the distal poly(A) site. If a splice site was identified, then the APA event was termed as alternative last exon (ALE), while poly(A) sites lacking an intervening splice site and hence located on the same last exon were termed tandem UTR (TUTR) events. In cases where either the proximal or distal poly(A) sites mapped to multiple transcript isoforms, some proximal-site isoforms may share a terminal exon with one of the distal-site isoforms, while others use a different last exon (and are thus separated by a splice site). As different pairs of transcripts can yield either a shared or an alternative last exon, these events were termed MIXED. When the proximal and distal poly(A) sites were assigned to the same transcript isoform but the proximal poly(A) site was located upstream of the beginning of the annotated 3’ UTR, then the APA event was termed internal APA (iAPA). The coordinates of 3’ UTR boundaries as well as the location of all splice sites were extracted from the gencode.v45.annotation.gtf annotation file using the GTFtools package^66^.

### Categorisation of DMAi mitigation status among siCFIM25-regulated genes

Firstly, genes that displayed a statistically significant, greater than 10% (+0.1) increase in proximal poly(A) site usage and a statistically significant, greater than 10% (-0.1) decrease in distal poly(A) site usage upon siCFIM25, such that the overall proximal to distal usage shift was at least less than -0.2, were classed as siCFIM25-shortened genes. Only TUTR genes were retained, to avoid splicing-related events. Then ratios between the proximal to distal shift displayed by these genes following siCFIM25 + DMSO treatment versus the proximal to distal shift triggered by siCFIM25 + DMAi treatment were calculated in order to designate DMAi mitigation status, according to the following logic:

Full DMAi mitigation = siCFIM25 + DMAi prox-to-dist shift <20% of siCFIM25 + DMSO prox-to-dist shift.
Partial DMAi mitigation = siCFIM25 + DMAi prox-to-dist shift between 30%-60% of siCFIM25 + DMSO prox-to-dist shift.
No DMAi mitigation = siCFIM25 + DMAi prox-to-dist shift between 80-120% of siCFIM25 + DMSO prox-to-dist shift.
DMAi exacerbated = siCFIM25 + DMAi prox-to-dist shift >133% of siCFIM25 + DMSO prox-to-dist shift.

### Categorisation of DMAi vs non-DMAi poly(A) site groups

To be classed as a common DMAi-regulated site, the poly(A) site firstly had to be present in at least two DMAi-treated cancer line datasets (after passing the DRIMSeq read coverage thresholds described above), with an average DMAi-induced change in usage across these lines of at least 10%, and a minimum change in usage of 7.5% in every line. This change in usage had to pass the significance threshold (adjusted p value < 0.05) in at least one of the lines. Our collated panel of non-DMAi datasets comprised several previously published 3’ RNA-seq datasets: CBPi^67^, CDK12i^68^, CLKi^69^, CPSF73i (JTE607)^41^, METTL3i^70^, siPCF11^40^, siSUPT4H1^71^ and siXRN2^72^, as well as numerous internally generated 3’ RNA-seq datasets: CDK11i/TOPKi, siCFIM25, siELAVL1 and siSam68. Poly(A) sites that had passed the above common DMAi filters were then only retained as common DMAi-regulated sites if the average DMAi-induced change in usage was at least 33% the maximum change in usage in a non-DMAi dataset. To be classed as a non-DMAi site, the poly(A) site had to display a change in usage of at least 10% with an adjusted p value < 0.05 in at least one non-DMAi dataset. Then, the average DMAi effect was required to be less than 33% of the non-DMAi induced change in usage. In the case of each group, the proximal and distal sites had to each pass the described filters for the gene to be classed as a DMAi or non-DMAi gene. Genes were classified as control genes if both poly(A) sites averaged a change in usage of less than 7.5% across DMAi and non-DMAi datasets. In each category, only TUTR APA genes were retained, to avoid splicing-related events.

### Motif frequency analysis

Discovery of enriched motifs was performed using STREME^73^ (v5.5.7) in RNA mode, using sequences -100 to +100 surrounding the poly(A) site. To subsequently evaluate the frequency of these motif groups surrounding poly(A) sites, we traversed each sequence recording whether the motifs were present or absent at each position, and aggregated the counts at each position across all sites. The results were visualized using a running average over windows of 10, 15, or 40 nucleotides (as defined by the motif group, specified in the text), with the window shifting by one nucleotide at a time.

### RBP eCLIP binding analysis

Raw ENCODE eCLIP fastq files for each RBP analysed in both HepG2 (103 RBPs) and K562 (120 RBPs) were downloaded from www.encodeproject.org, and processed using the nf-core/clipseq^74^ (v1.0.0) pipeline to generate crosslink bed files, as described in Kuret et al., 2022^75^. Individual replicates were merged using BEDtools. These merged crosslink files were then read into R as GRanges^76^ objects, and resized to a 400-nt window centered on the poly(A) site (200 nt upstream/downstream). Using the ScoreMatrix function from the genomation package^76,77^, crosslink coverage was computed and summarized into mean coverage meta-profiles for each condition.

## CODE AVAILABILITY

The code used to analyse the data and to produce the figures in this work is available on Github (https://github.com/ulelab/PRMT-APA/tree/main/Scripts/scripts_for_figures_in_paper/) and is also archived on Zenodo (DOI: 10.5281/zenodo.14968061).

---

**Figure S1.**
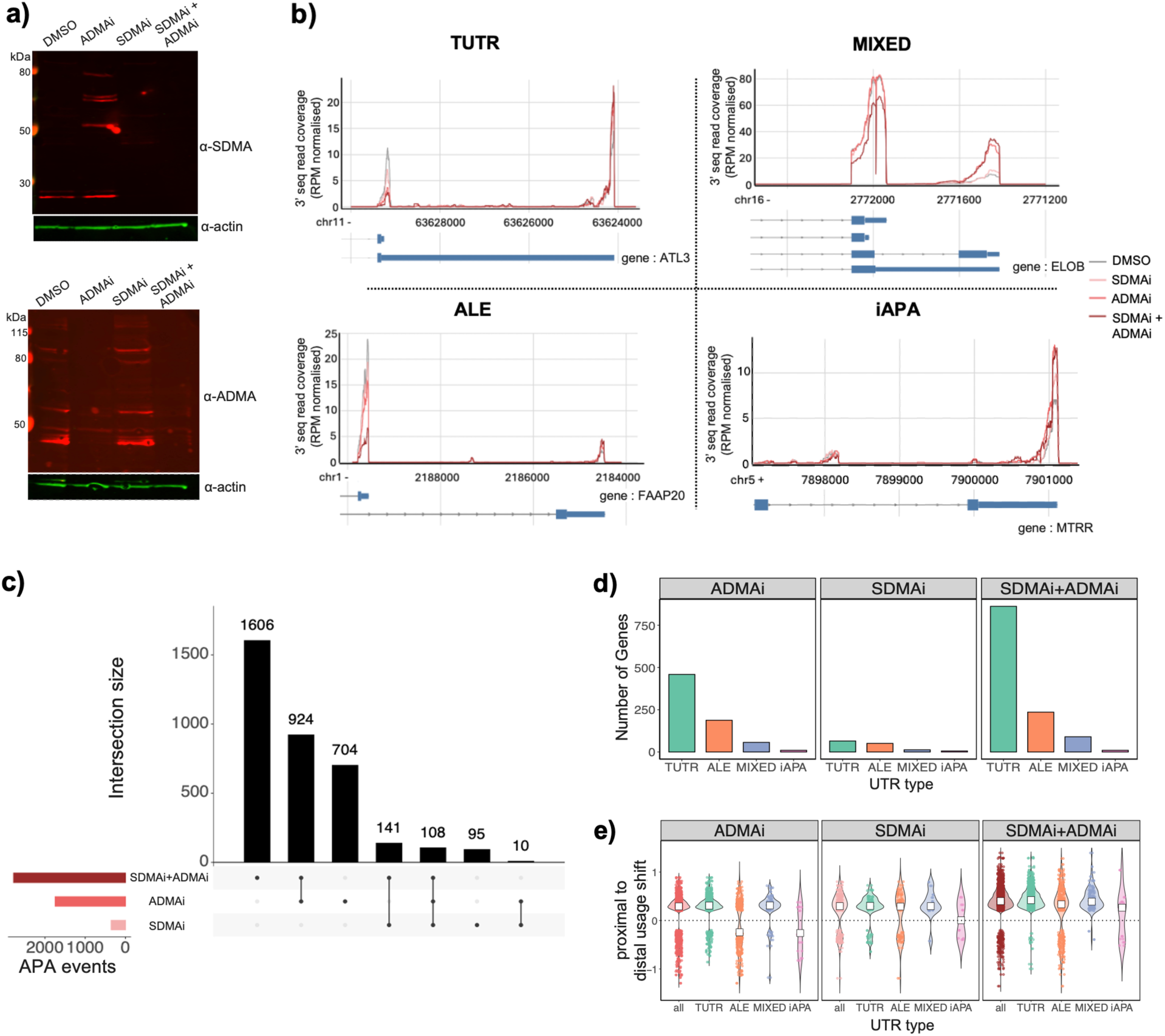
**a)** Western blot showing the reduction in SDMA and ADMA in LU-99 cells upon 96 hours treatment with GSK-715 (ADMAi), LLY-283 (SDMAi), or both (SDMAi + ADMAi). **b)** Examples of a same terminal exon APA event (TUTR) (Upper left), an alternate last exon APA event (ALE) (Bottom left), an APA event that can involve either the same terminal or an alternate last exon (MIXED) (Upper right), and an APA event involving an internal exon or intron (iAPA) (Bottom right). **c)** Upset plot visualising the number of significant APA events triggered by SDMAi, ADMAi or SDMAi+ADMAi and their intersections. poly(A) sites with a greater than 10% usage change and adjusted p value < 0.05 were deemed significant APA events. **d)** Bar chart displaying the number of significant APA events belonging to each APA class following arginine methylation inhibition. **e)** Violin plot showing proximal to distal poly(A) site usage shifts among each class of significant APA event. Proximal to distal usage shift is calculated as: dPA change in usage - pPA change in usage. White square = median.

**Figure S2.**
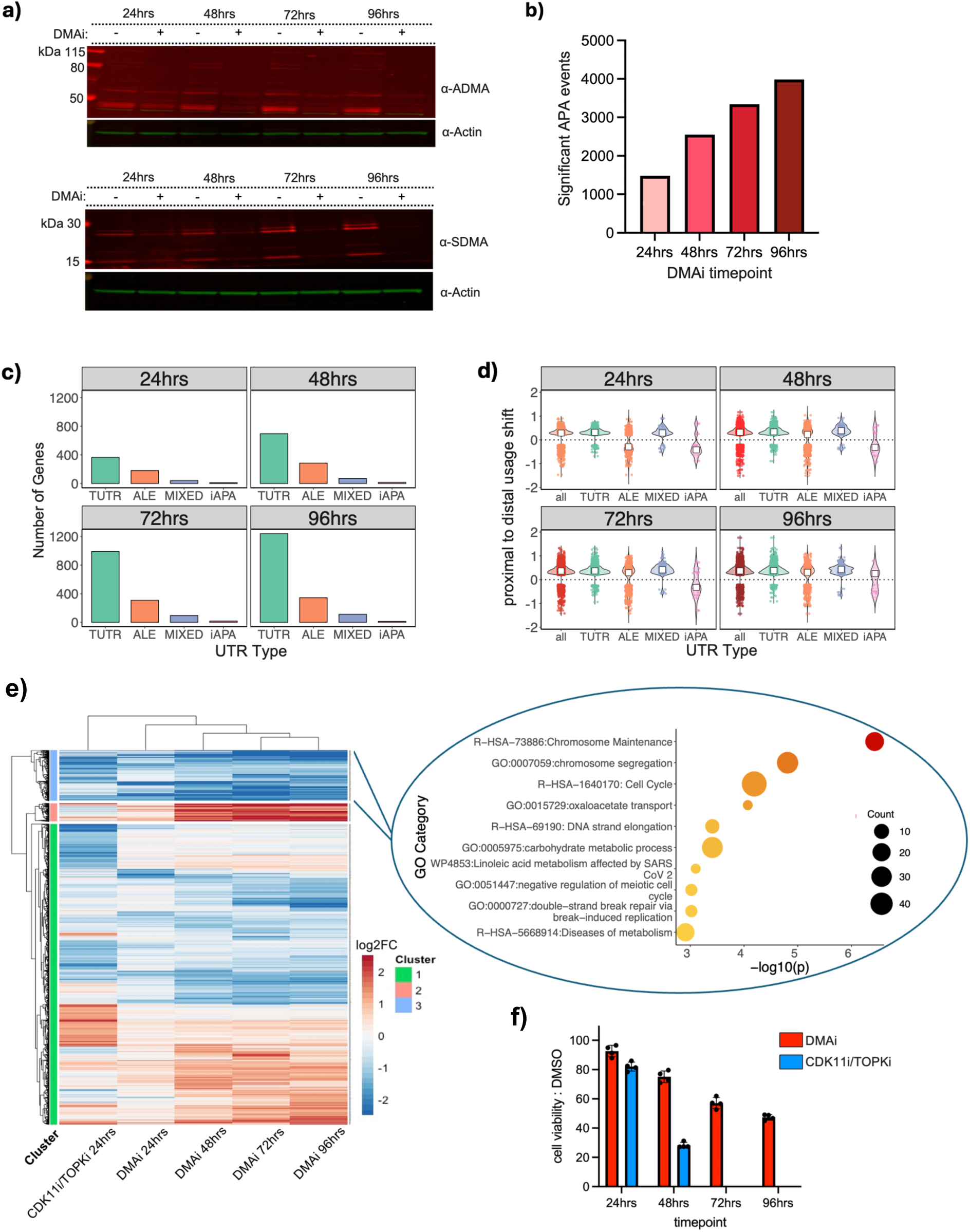
**a)** Western blot demonstrating the reduction in SDMA and ADMA in LU-99 cells upon treatment with GSK-715 (ADMAi) and LLY-283 (SDMAi) for 24, 48, 72 or 96 hours. **b)** Bar chart outlining the number of significant APA events induced by DMAi at each time point. poly(A) sites with a greater than 0.1 (10%) usage change, adjusted p value < 0.05, were deemed significant APA events. **c)** Bar chart showing the number of significant APA events belonging to each class of APA at each time point. **d)** Violin plot showing proximal to distal poly(A) site usage shifts among each class of significant APA event. Proximal to distal usage shift is calculated as: dPA change in usage - pPA change in usage. White square = median. **e)** Left - Heatmap displaying significant log2Foldchange in gene expression in LU-99 cells following CDK11i/TOPKi or DMAi. Right - Gene ontology term enrichment analysis of genes located in Cluster 3 in heatmap. **f)** Bar chart displaying LU-99 cell viability at 24 hour intervals following treatment with either GSK-715 + LLY-283 (SDMAi + ADMAi) or OTS964. Cell viability was determined using the CellTiter-Glo® 2.0 Cell Viability Assay, and is expressed relative to cells treated with DMSO.

**Figure S3.**
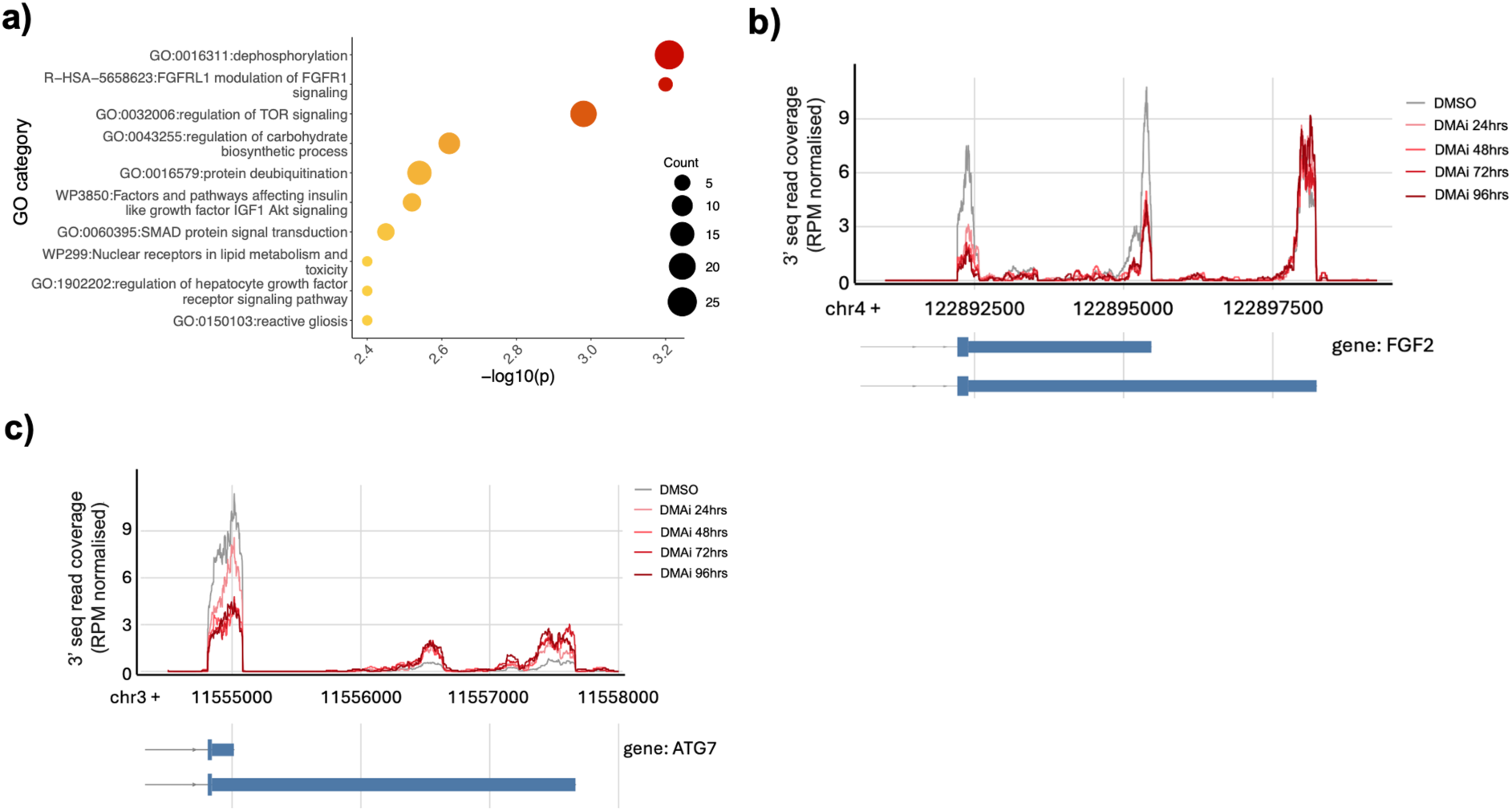
**a)** Gene ontology term enrichment analysis of genes displaying significant APA upon 24 hours of DMAi treatment. **b)** Visualisation of change in FGF2 pPA and dPA site usage in LU-99 cells triggered by DMAi. **c)** Visualisation of change in ATG7 pPA and dPA site usage in LU-99 cells triggered by DMAi.

**Figure S4.**
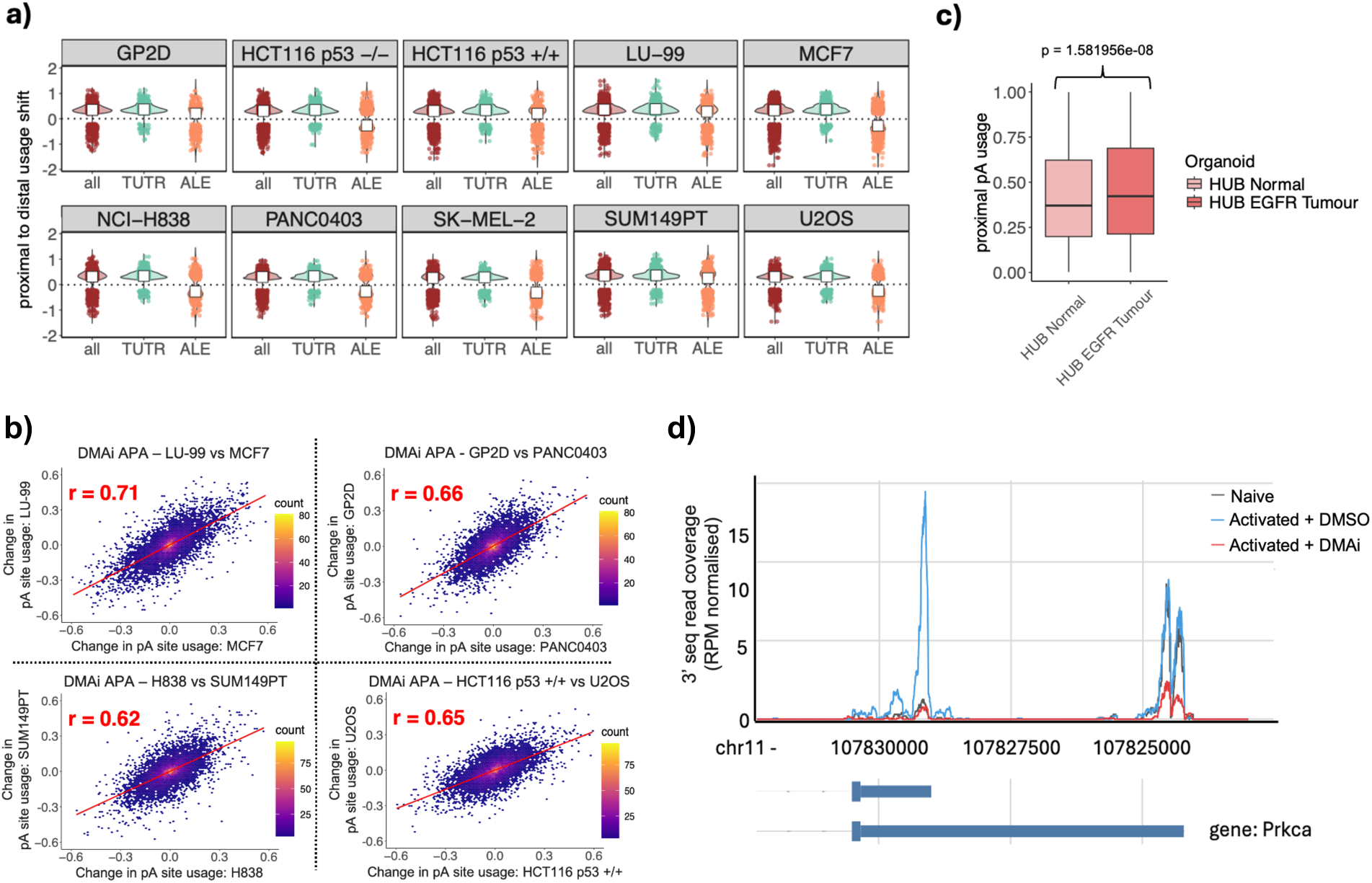
**a)** Violin plot showing proximal to distal usage shifts across each class of significant APA event. Proximal to distal usage shift is calculated as: dPA change in usage - pPA change in usage. White square = median. **b)** Scatter plots displaying Pearson correlation (r) of DMAi-induced poly(A) site usage change between different cell lines. Upper left = LU-99 vs MCF7, Lower left = NCI-H838 vs SUM149PT, Upper right = PANC0403 vs GP2D. Lower right = HCT116 p53 +/+ vs U2OS. **c)** Distribution of proximal poly(A) usage across all genes in a patient-derived lung tumour organoid model versus normal lung tissue organoids derived from the same patient. Scale: 0 = 0% usage, 1 = 100% usage. Statistical significance was calculated using the Kruskal-Wallis Test. **d)** An example of a 3’ UTR (*Prkca)* whose shortening upon T cell activation (blue) is prevented by DMAi (red).

**Figure S5.**
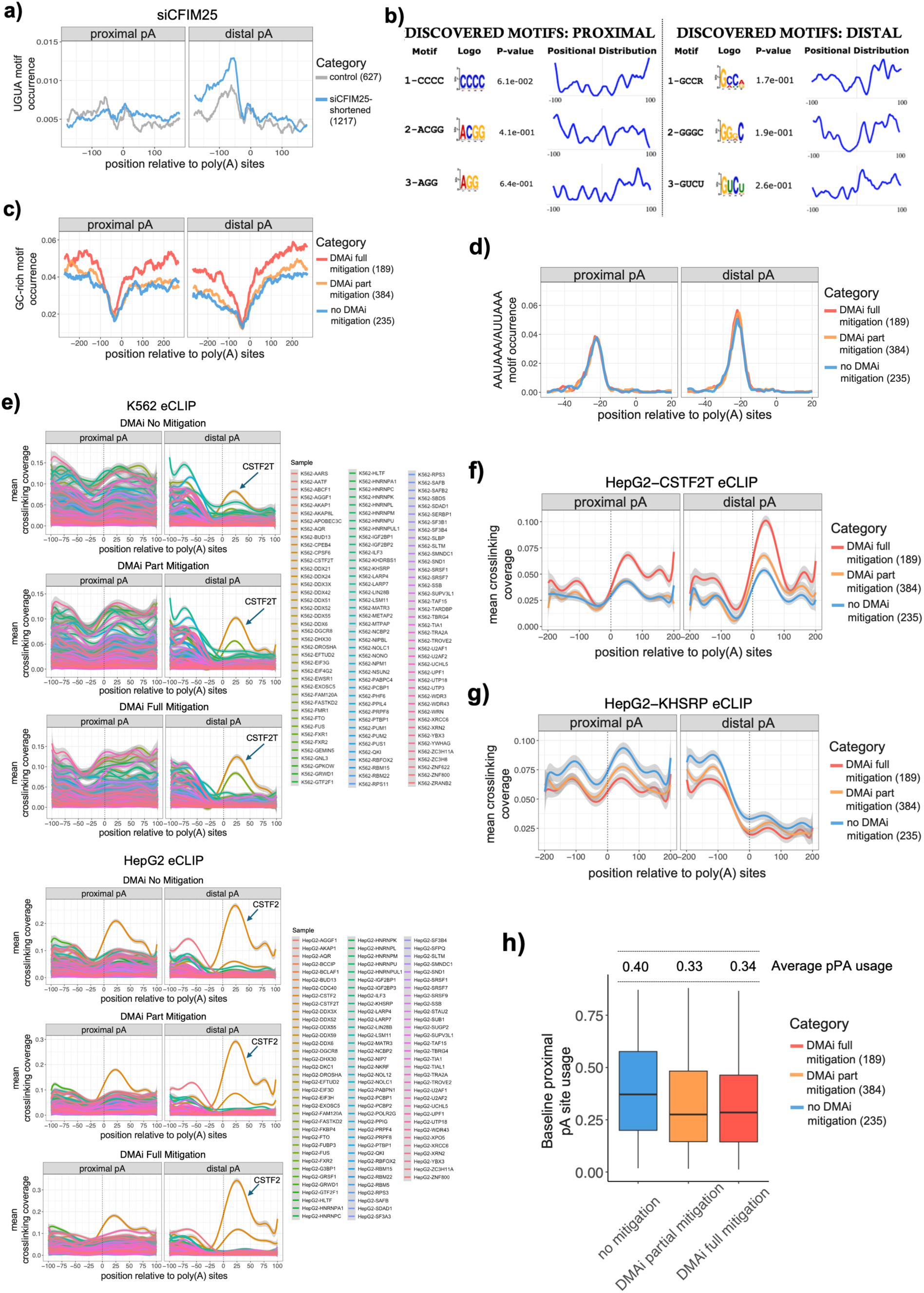
**a)** Comparison of position-dependent frequency of occurrence of UGUA within proximal and distal poly(A) site regions at siCFIM25-shortened 3’ UTRs. Running averages were calculated over 30 consecutive nucleotide positions. **b)** STREME motif enrichment results displaying the most enriched motifs identified at proximal pA regions (left) and distal pA regions (right) within DMAi-full mitigation versus DMAi-no mitigation CFIM25-regulated transcripts. **c)** Comparison of position-dependent frequency of occurrence of GC-rich motifs within proximal and distal regions at DMAi-full mitigation (red), partial mitigation (orange) and no mitigation (blue) siCFIM25-shortened 3’ UTRs. Running averages were calculated over 40 consecutive nucleotide positions. GC-rich motif = CCC, GGG, or any 3mer combination of 2Gs + 1C / 2Cs + 1G. **d)** Comparison of position-dependent frequency of occurrence of AAUAAA or AUUAAA within proximal and distal regions in 3’ UTRs across each DMAi sensitivity category. Running averages were calculated over 10 consecutive nucleotide positions. **e)** Upper = RBP eCLIP signal in K562 cells across each DMAi sensitivity category (expression matched), calculated from ENCODE consortium data. Lower = RBP eCLIP signal in HepG2 cells across each DMAi sensitivity category (expression matched), calculated from ENCODE consortium data. **f)** CSTF2T eCLIP binding around poly(A) sites in 3’ UTRs across each DMAi sensitivity category (expression matched), calculated from ENCODE consortium data in HepG2. **g)** KHRSP eCLIP binding around poly(A) sites in 3’ UTRs across each DMAi sensitivity category (expression matched), calculated from ENCODE consortium data in HepG2 cells. **h)** Distribution of baseline proximal poly(A) usage across each DMAi sensitivity category (expression matched). Scale: 0 = 0% usage, 1 = 100% usage.

**Figure S6.**
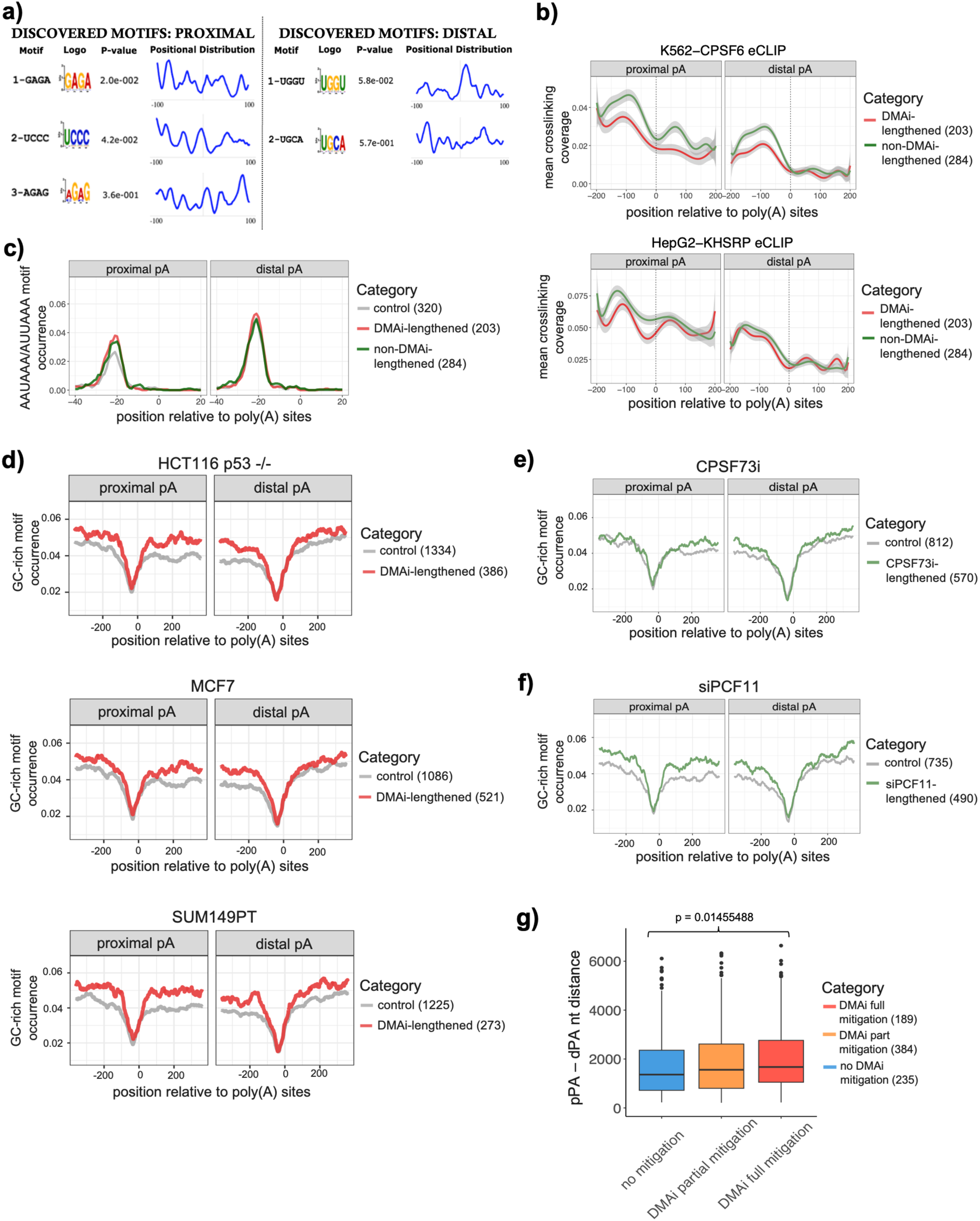
**a)** STREME motif enrichment results displaying the most enriched motifs identified within DMAi-lengthened 3’ UTRs at proximal and distal regions compared to non-DMAi-lengthened 3’ UTRs. **b)** eCLIP crosslinking profiles (obtained from ENCODE consortium data) around poly(A) sites in 3’ UTRs of DMAi or expression matched non-DMAi-lengthened transcripts, calculated from ENCODE consortium data. Upper plot shows CPSF6 data from K562 cells, and lower plot the KHRSP data from HepG2 cells. **c)** Comparison of position-dependent frequency of occurrence of AAUAAA or AUUAAA within proximal and distal regions at DMAi-lengthened (red), non-DMAi-lengthened (green) and control (grey) sites. Running averages were calculated over 10 consecutive nucleotide positions. **d)** Comparison of position-dependent frequency of occurrence of GC-rich motifs within proximal and distal regions at DMAi-lengthened (red) and unchanged, control (grey) 3’ UTRs, across three representative cancer lines. Running averages were calculated over 40 consecutive nucleotide positions. Upper = HCT116 p53 -/-, Middle = MCF7, Lower = SUM149PT. GC-rich motif = CCC, GGG, or any 3mer combination of 2Gs + 1C / 2Cs + 1G. **e)** Comparison of position-dependent frequency of occurrence of GC-rich motifs within proximal and distal regions at JTE607 (green) and unchanged, control (grey) 3’ UTRs in HepG2 cells. Running averages were calculated over 40 consecutive nucleotide positions. JTE607 3’ RNA-seq libraries were previously generated by Cui et al.^41^, and reprocessed using our APA pipeline. **f)** Comparison of position-dependent frequency of occurrence of GC-rich motifs within proximal and distal regions at siPCF11 (green) and unchanged, control (grey) 3’ UTRs in HeLa cells. Running averages were calculated over 40 consecutive nucleotide positions. siPCF11 3’ RNA-seq libraries were previously generated by Kamieniarz-Gdula et al.^40^, and reprocessed using our APA pipeline. **g)** Boxplot showing the nucleotide distance between proximal and distal poly(A) sites at siCFIM25-shortened 3’ UTRs that are not sensitive to DMAi (no mitigation = blue), partially sensitive (DMAi partial mitigation = orange), and fully sensitive (DMAi full mitigation = red).

